# Environmental Frass Remodels the Larval Gut Microbiome to Enable Black Soldier Fly (Hermetia illucens) Bioconversion of Livestock Manure

**DOI:** 10.64898/2026.11.05.764513

**Authors:** Audrey Pettengill, Rachel Stender, Kendyl Thitathan, Michael Cleary

**Affiliations:** Department of Molecular and Cell Biology, University of California, Merced, Merced, USA

## Abstract

Black Soldier Fly larvae (BSFL) consume nearly any organic material and show promise for the remediation of agricultural waste, including livestock manure. BSFL feeding generates frass, a mixture of unconsumed substrate, larval feces, and microorganisms. We identified a wild population of BSFL (Yuba strain) thriving on rabbit manure, a challenging substrate due to its high lignocellulose content. We investigated if frass extract from this source was necessary and sufficient to support bioconversion of manure and if such effects are altered by captive husbandry. Larvae supplemented with domestic frass failed to grow on rabbit manure but grew well when supplemented with wild-sourced frass. Microbiome analysis revealed that wild-sourced frass contributes a cohort of lignocellulose-degrading microorganisms to the BSFL gut, particularly *Rummeliibacillus* bacteria and *Scedosporium* fungi. These growth-promoting effects extended to cattle manure. Following initial seeding with wild-sourced frass, lab-cultured frass ultimately failed to support growth on rabbit manure and the gut microbiome shifted toward bacteria commonly found in domestic BSFL. This study reveals that frass extract derived from environmental sources can improve the bioconversion ability of BSFL and that beneficial microbes are lost during captive-rearing.

**Importance:** Livestock manure is an abundant source of environmental pollution. Black Soldier Flies have the potential to bioconvert manures and mitigate associated harms. Many studies have investigated the role of substrate and larval gut microbiomes in bioconversion but have ignored microbiomes from wild BSFL microecosystems. This work investigates whether frass extract from a wild population is sufficient to remodel the BSFL gut microbiome and enable growth on lignocellulose-rich animal manures. The focus on wild frass revealed novel determinants of BSFL growth and our results may inform development of waste bioconversion systems optimized by targeted frass manipulations.

---

Global livestock production has reached record-high levels and is projected to increase alongside human population growth and societal changes in diet preferences^1^. Based on current animal agriculture practices, this growth will be accompanied by environmental impacts, including greenhouse gas emissions and pollution of aquatic ecosystems^2–4^. The drivers of these impacts are multifaceted, but one significant contributor is livestock manure. Manure accounts for 10% of total greenhouse gas emissions from livestock^4^ and methane production from microbial decomposition of cattle manure may be a greater contributor to global warming than enteric methane emissions^5^. Waterways are harmed when nitrogen and phosphorous-rich manure runoff causes eutrophication^6^. Many strategies exist for managing livestock manure^7^, including composting systems that reduce pollution while converting the waste into valuable products like fertilizer or biogas. One approach involves feeding manure to larvae of the Black Soldier Fly (*Hermetia illucens*)^8^; the manure is bioconverted into insect mass and the resulting Black Soldier Fly larvae (BSFL) are a valuable source of biomaterials^9^. This waste reduction and valorization scheme can support a circular system in which BSF produced via manure consumption are employed in subsequent rounds of manure bioconversion.

The efficiency of waste bioconversion depends on an intersection of fly genetics^10–14^ and environmental factors^15^. Among the most influential environmental factors are the microbial communities of the substrate and the larval gut^16–21^. The mixture of substrate, BSFL feces, shed larval cuticles, and microorganisms is known as frass. A complex relationship exists between frass and larval gut microbiomes; host selection partially determines which microbes colonize the gut and larval feces may in turn modify the frass microbiome^22–24^. Lignocellulose-rich organic wastes, such as rabbit and cattle manure, present an especially challenging substrates for BSFL^8,23,25–28^, mainly because the fly genome does not encode enzymes to catabolize cellulose, hemicellulose or lignin^29^. The solution to this challenge comes from frass and larval gut microorganisms that are able to degrade lignocellulose^28,30^, but how such metabolic functions are established and maintained within BSF populations is not well understood. While previous work has provided insights into how frass and gut microbiomes allow BSFL to bioconvert organic wastes, these studies typically use domestic fly strains and naïve (BSFL-free) substrates^17,31,32^. Similarly, experiments using wild BSF typically provide larvae with naïve substrates, free of frass from the relevant natural habitat^33–35^. Such approaches overlook wild BSFL microecosystems that have been optimized by natural selection. Very little is known about the ecological dynamics of wild BSF, despite native and naturalized populations occuring across much of the globe, in niches ranging from carrion to honeybee colonies^36–38^. BSFL microecosystems represent a vast untapped resource for discovering aspects of fly behavior, genetics and microbial symbiosis that could be co-opted to improve waste remediation applications^39^. Wild BSFL gut microbiomes are more diverse than domestic gut microbiomes and vary by geographic location^40^. Environmental frass is potentially equally diverse and may harbor microorganisms that support BSFL bioconversion of manure and other substrates.

We discovered a wild population of BSF in northern California that was thriving on rabbit manure, a challenging lignocellulose-rich substrate. This microecosystem was recapitulated in the laboratory and used to test if environmental frass-associated microbes were necessary and sufficient to support growth on lignocellulose-rich animal manure. This system also allowed investigation of how captive rearing alters frass-dependent effects on BSFL growth and microbiome composition.

## RESULTS

### Wild and domestic *Hermetia illucens* populations are genetically similar

Rabbit manure containing BSFL was collected from a farm in Yuba County, California, USA. Naïve rabbit manure without any BSFL was collected in parallel from the same location. Adult flies emerged directly from the rabbit manure sample within laboratory cages, and this population was captive bred for six generations. We used established microsatellite markers^36^ to assess population genetic differences among the Yuba P0 (Y.P0) generation, the Yuba F6 (Y.F6) generation, and a commercial strain derived from North American BSF domesticated over 25 years ago^41^, here referred to as the Domestic strain. Principle component analysis placed each population in distinct but neighboring distributions, with Y.F6 and Domestic nested within the broader Y.P0 distribution (Fig. 1A). Inbreeding coefficients and allelic richness were not significantly different between Y.P0 and Y.P6 while the Y.P0 population was significantly less inbred and more allele rich than the Domestic population (Fig. 1B). Two measures of genetic differentiation, the fixation index (F_ST_) and Jost’s D, revealed only minor genetic distances between these populations (Fig. 1C). This genetic information was used in the design and interpretation of larval growth and microbiome experiments described below.

**Figure 1.**
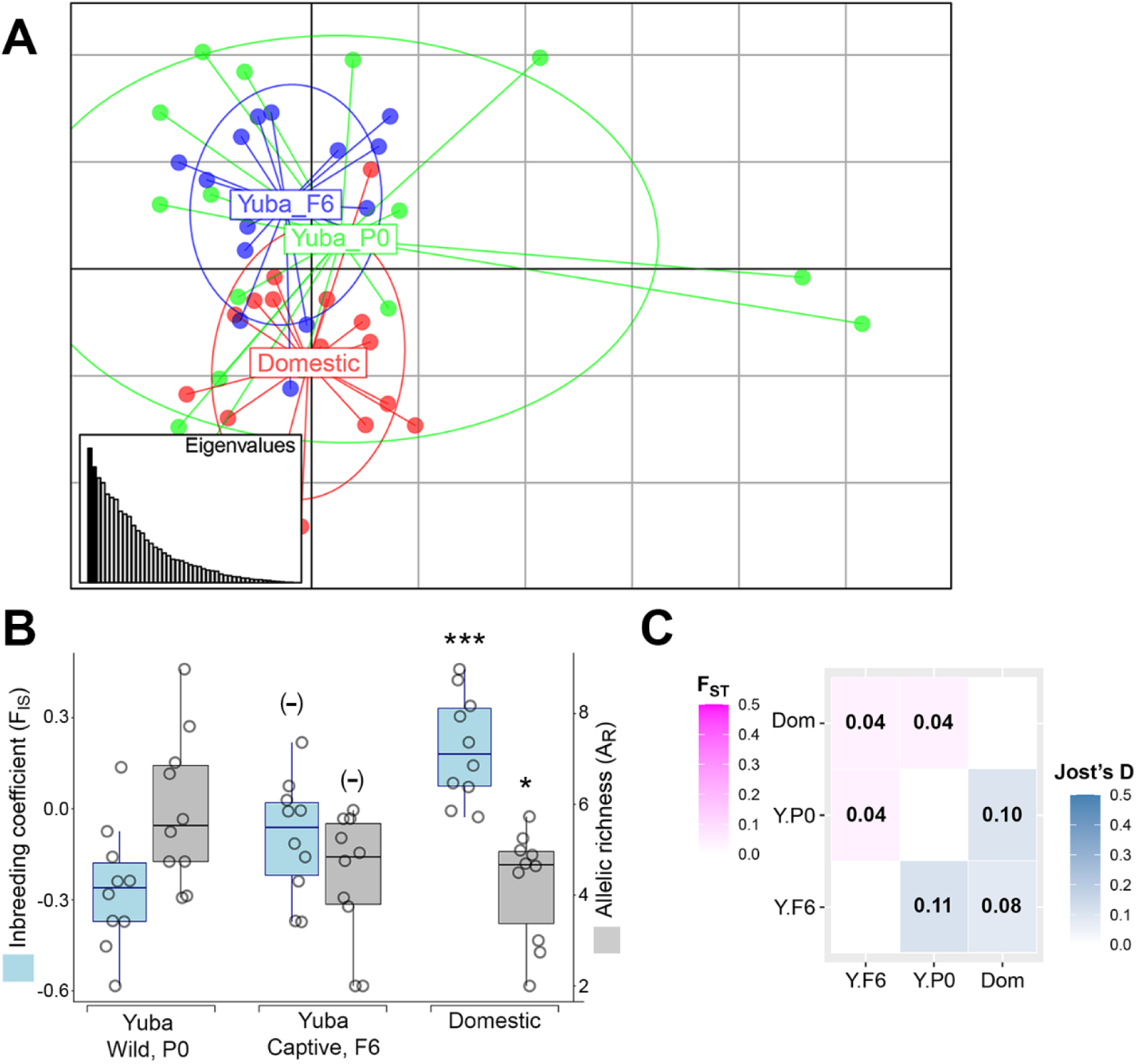
Yuba and domestic populations are genetically similar. **A**. Principle component analysis of microsatellite data. Eigenvalues corresponding to represented components are filled black. Points represent genotypes; populations are color-coded and labeled within their 95% inertia ellipses, 16 individuals per population. **B**. Distribution of inbreeding coefficient (F_IS_) and allelic richness (A_R_) for ten microsatellite loci in each population. (-) indicates no significant difference compared to Yuba Wild, P0. *** indicates p-value < 0.0001, * indicates p-value = 0.05 compared to Yuba Wild, P0. **C**. Genetic differentiation among populations as measured by the fixation index (F_ST_) and Jost’s D. Color scales represent a range from no differentiation to high differentiation. Values for each pairwise comparison are given.

### Wild-sourced frass extract is necessary and sufficient for growth on rabbit manure

Yuba and Domestic populations were used to test how frass extracts affect growth on rabbit manure. Frass extract was prepared by filtering total frass through a 660 µm mesh and liquid extract was inoculated into naïve rabbit manure at 5% concentration. Frass samples were: “wild-sourced frass” = the original farm-collected rabbit manure + BSFL after being established in laboratory cages, “domestic frass” = lab diet + Domestic BSFL, and “captive frass” = lab diet + captive-bred Yuba at the indicated generation. Captive frass samples were seeded with frass from the preceding generation via larval feeding and defecation (see Methods). Based on this design, we predicted that microbes initially derived from wild-sourced frass would be sequentially diluted, with highest abundance in the F1 captive frass (lab diet seeded by P0 larvae) and lowest abundance in the F5 captive frass (lab diet seeded by F4 larvae).

We began testing growth on rabbit manure at the Yuba F2 generation. Eggs were directly hatched onto the substrate in all experiments. In the absence of frass extract, Y.F2 larvae grew very poorly (Fig. 2A). Inoculation with domestic frass extract failed to augment growth but inoculation with wild-sourced frass extract significantly increased larval growth (Fig. 2A). This growth-promoting effect was lost if wild-sourced frass extract was sterile-filtered, indicating that growth depends on microbes in the extract (Fig. 2A). Inoculation with captive frass extract revealed that growth-promoting effects are lost during the captive rearing process. Y.F2 larvae supplemented with F1 captive frass grew only slightly worse than those supplemented with wild-sourced frass, but Y.F6 larvae supplemented with F5 captive frass grew no better than those supplemented with domestic frass (Fig. 2A). To determine if wild-sourced frass extract supports growth regardless of BSF strain, we analyzed growth of Domestic larvae on rabbit manure. As observed for the Yuba strain, only wild-sourced frass extract supported growth (Fig. 2B).

**Figure 2.**
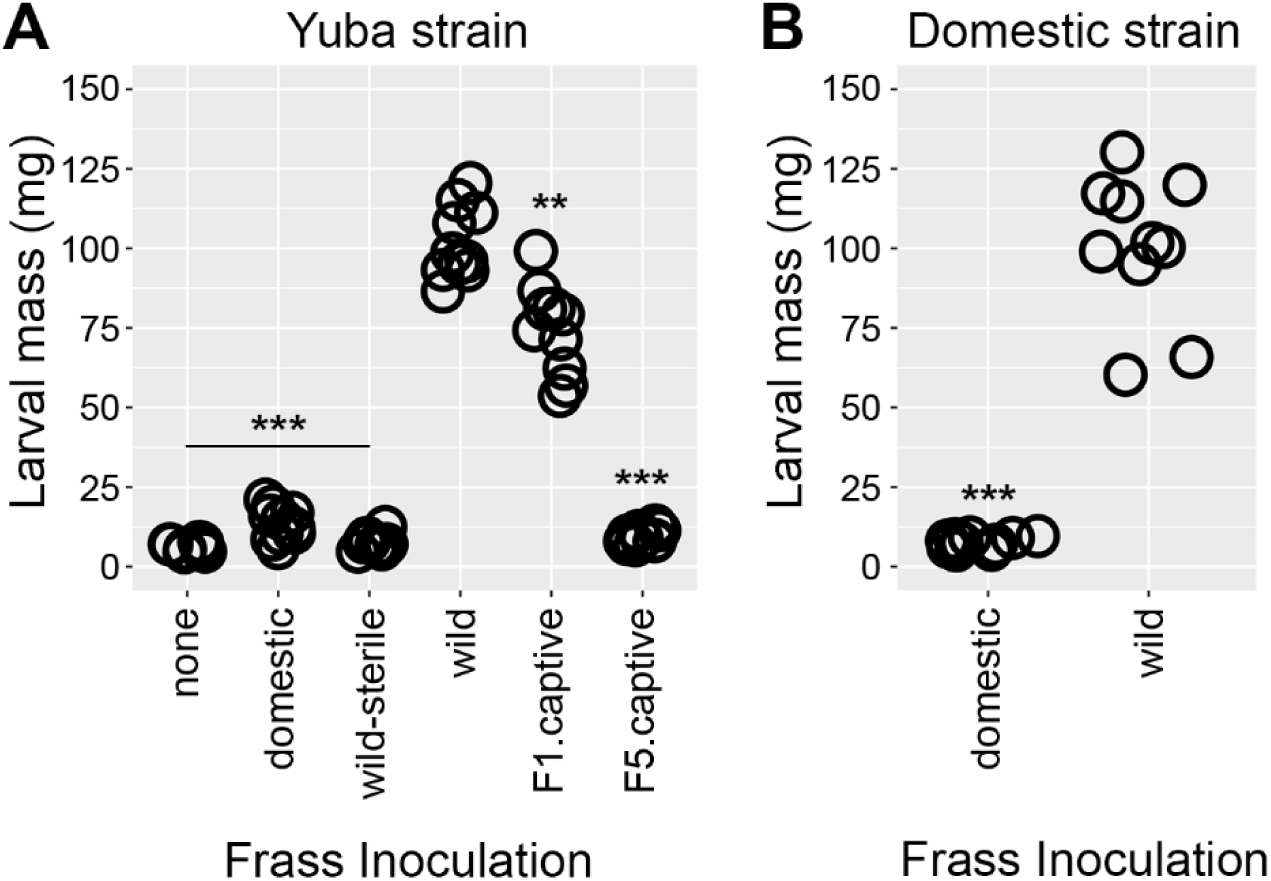
Wild-sourced frass extract enables growth on rabbit manure. **A**. Yuba strain larval mass measurements after rearing on rabbit manure for 21 days. n = 10 larvae per condition. Frass treatment is indicated on the x-axis. F5.captive frass extract was applied to Yuba F6 larvae, all other experiments used Yuba F2 larvae. *** indicates p-value < 0.0001, ** indicates p-value < 0.001 compared to wild frass treated. Wild-sterile is wild frass extract after passage through a 0.2 µm filter. **B**. Domestic strain larval mass measurements after rearing on rabbit manure for 21 days. n = 10 larvae per condition. Frass treatment is indicated on the x-axis. *** indicates p-value < 0.0001 compared to wild frass treated.

### A small, reproducible cohort of bacteria and fungi are linked to frass-dependent growth

To identify microbes that may explain frass-dependent growth on rabbit manure, we performed bacterial and fungal microbiome sequencing of substrate frass and BSFL guts. Comparisons of operational taxonomic units (OTUs) across biological replicates and varying treatments revealed dynamic changes in microbial communities (Fig. 3). While substrate frass OTU abundance was largely uniform across treatments and failed to correlate with growth phenotypes, there were several gut OTU patterns of interest, particularly those with higher levels in the gut following wild-sourced frass treatment compared to domestic frass treatment. Five bacterial OTUs were significantly enriched in the gut of Y.F2 larvae treated with wild-sourced frass. These are OTUs that map to the following (listed in descending order of enrichment): genus *Rummeliibacillus*, family *Porphyromonadaceae*, genus *Bacillus*, genus *Paenibacillus*, and genus *Dysgonomonas* (Fig. 4A). Two fungal OTUs were significantly enriched in the gut of Y.F2 larvae treated with wild-sourced frass: one mapped to the genus *Scedosporium* and one mapped to the genus *Petriella* (Fig. 4A).

**Figure 3.**
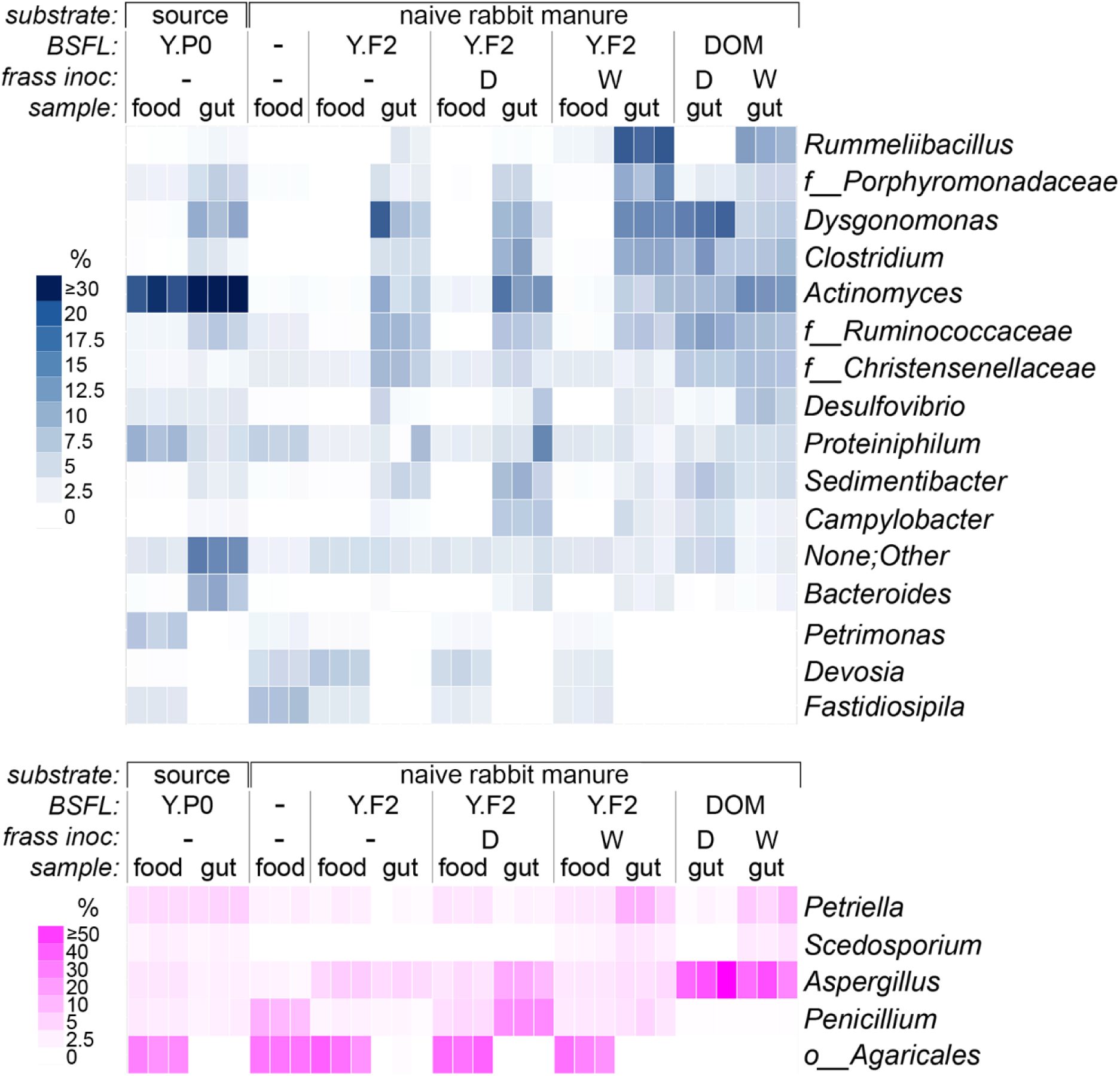
Substrate and gut microbiomes change in response to frass extract. Heatmap summary of bacterial (top, blue) and fungal (bottom, magenta) OTU abundance in food (substrate frass) and larval gut samples. Three biological replicates are shown per condition. Substrate frass identify is listed at the top: source = wild-sourced rabbit manure + BSFL, naive rabbit manure = BSFL-free rabbit manure. BSFL indicates the strain used or absence of larvae (-). Frass inoc. indicates the frass extract used: (-) = none, D = domestic, W = wild-sourced. OTU identities listed on the right refer to genus unless labeled f_ (family) or o_ (order). The heatmap color scale represents percent abundance for sequences mapping to the corresponding OTU. Only OTUs with an average abundance of ≥ 5% in at least one of the eleven conditions tested are included.

**Figure 4.**
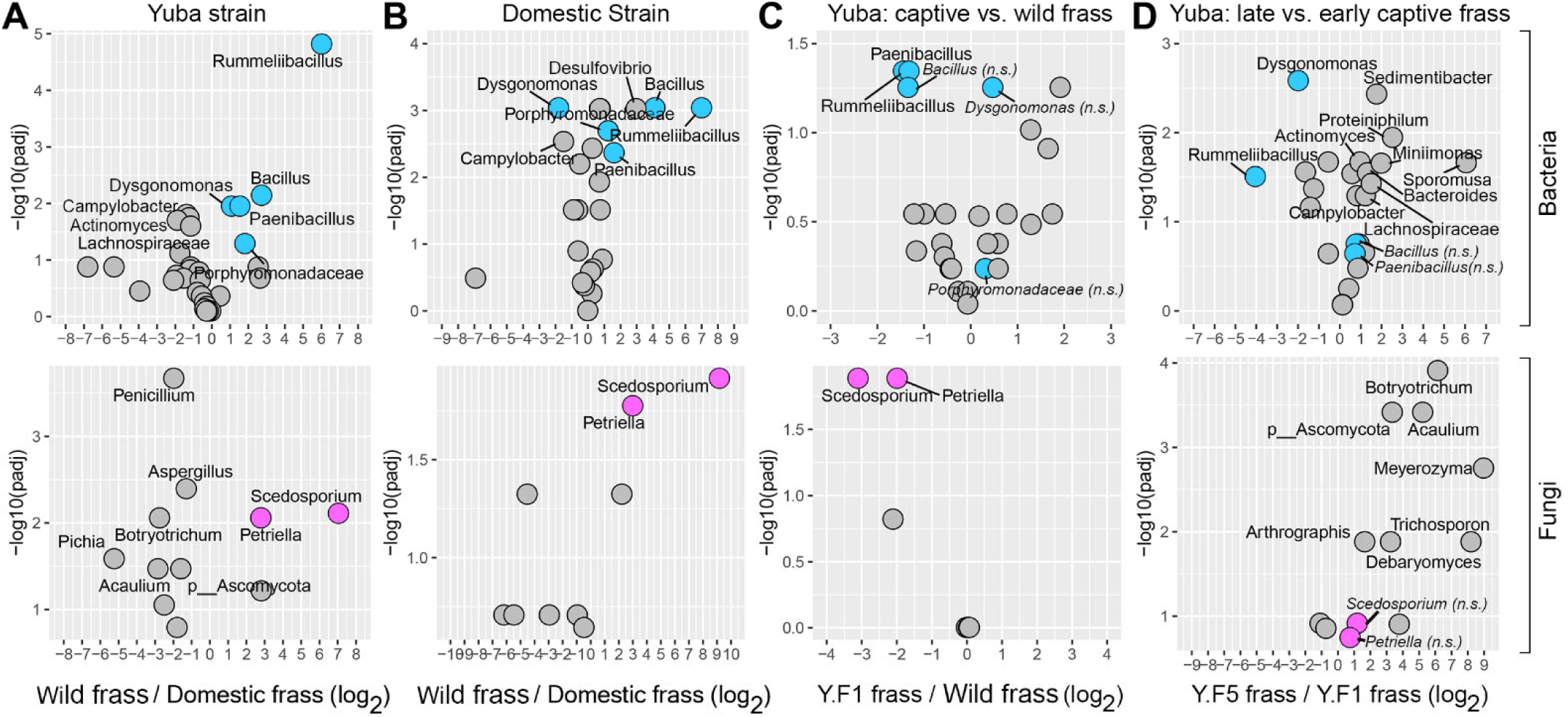
A small cohort of bacteria and fungi are enriched in wild frass-treated guts and depleted in captive frass treated guts. Volcano plots displaying fold-change in abundance (x-axis) and statistical significance (y-axis) for gut microbiome OTUs in the pairwise comparison listed at the bottom of each vertical panel. Bacterial data on top, fungal data on bottom. Bacteria of interest are labeled blue. Fungi of interest are labeled magenta. Only OTUs with fold-change greater than or equal to 2 (log_2_ ≥ 1.0) and adjusted p-values less than 0.05 (-log10(padj) > 1.30) are labeled by name. OTUs of interest that did not meet the adjusted p-value cutoff are labeled in italics and “n.s.” for not significant. **A**. Wild-sourced frass treatment OTU abundance divided by domestic frass treatment OTU abundance for Y.F2 larvae fed naive rabbit manure. **B**. Wild-sourced frass treatment OTU abundance divided by domestic frass treatment OTU abundance for Domestic larvae fed naive rabbit manure. **C**. Y.F1 captive frass treatment OTU abundance divided by wild-sourced frass treatment OTU abundance for Y.F2 larvae fed naive rabbit manure. **D**. Y.F5 captive frass treatment OTU abundance divided by Y.F1 captive frass treatment OTU abundance.

The same bacteria and fungi were selectively enriched in Domestic strain larvae that grew on rabbit manure, the only difference was a relative depletion of *Dysgonomonas* (Fig. 4B). The absence of *Dysgonomonas* enrichment in the Domestic strain narrowed the list of candidate microbes necessary for growth on rabbit manure to OTUs within genus *Rummeliibacillus*, family *Porphyromonadaceae*, genus *Bacillus*, and genus *Paenibacillus* plus fungal OTUs within the genus *Scedosporium* and genus *Petriella.* If these OTUs are necessary for growth on rabbit manure, their abundance may be expected to decrease in the gut of captive frass-treated BSFL. This is the general pattern we observed. Y.F2 larvae treated with F1 captive frass had significant decreases in *Rummeliibacillus*, *Paenibacillus*, *Scedosporium* and *Petriella* and equally low abundance of the other OTUs of interest (Fig. 4C). Similarly, Y.F6 larvae treated with F5 captive frass had significant decreases in *Rummeliibacillus*, *Scedosporium* and *Petriella* and equally low abundance of the other OTUs of interest (Fig. 4D).

### Captive rearing shifts the frass-dependent microbiome toward a set of commonly observed domestic gut bacteria

Given that wild-sourced frass promotes growth on nutrient-poor, difficult to digest food, we asked if the converse is true: does wild-sourced frass inhibit growth on nutrient-rich, easily digested lab diet? In the absence of frass inoculation, Yuba and Domestic larvae had relatively limited growth compared to frass-treated larvae (Fig. 5A), indicating a function for frass-derived microbes even on nutrient-rich food. Wild-sourced frass extract and domestic frass extract equally augmented growth on lab diet, independent of BSF strain (Fig. 5A). These results rule out an inhibitory effect of wild-sourced frass extract when larvae are grown on lab diet.

**Figure 5.**
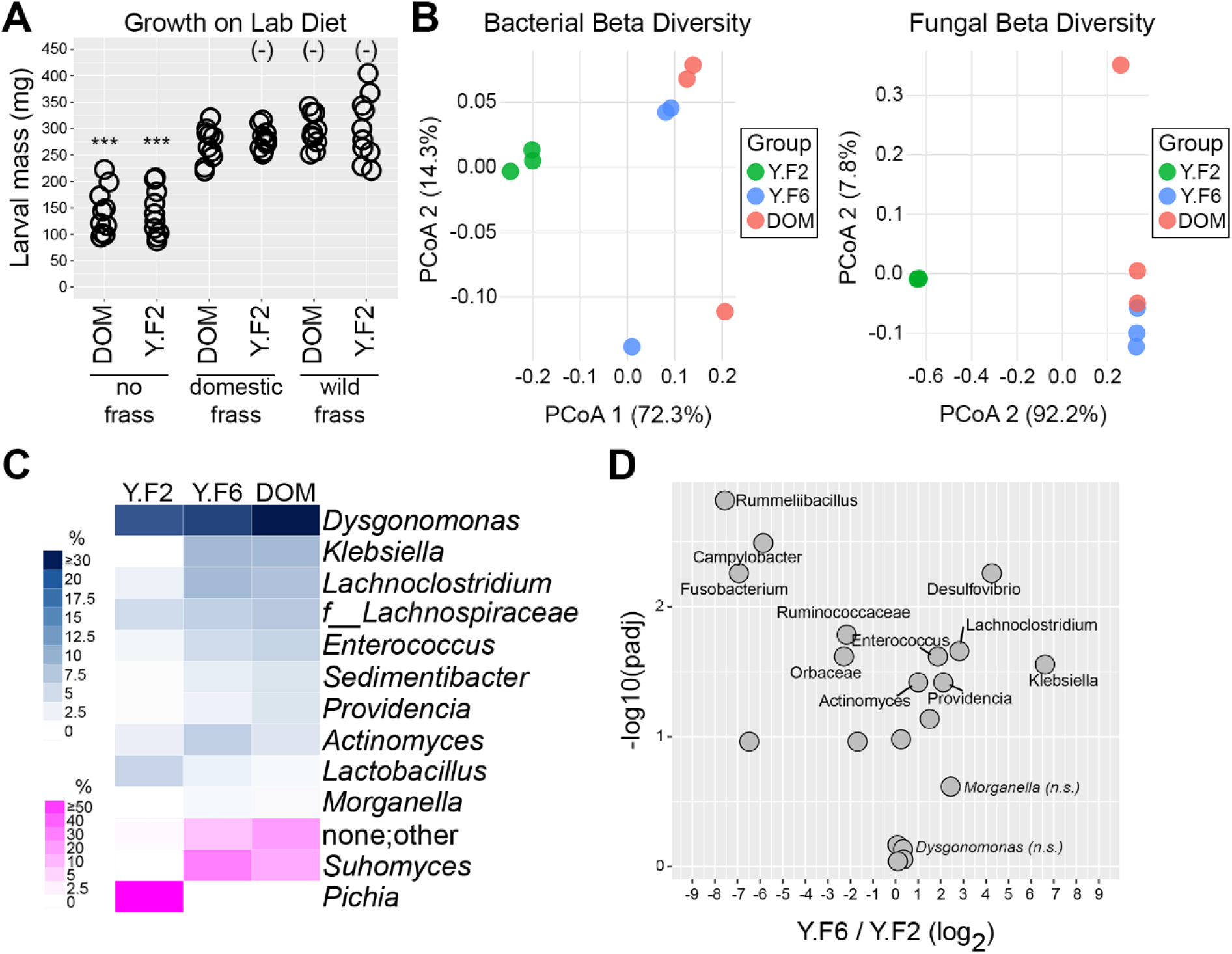
Captive rearing shifts the gut microbiome toward bacteria commonly found in domestic BSFL. **A**. Larval mass following 21 days of rearing on lab diet is shown for the strains and treatments indicated on the x-axis. DOM = domestic strain, Y.F2 = F2 Yuba. (-) indicates not significantly different from DOM larvae treated with domestic frass. *** indicates p-value < 0.0001 compared to DOM larvae treated with domestic frass. n = 10 larvae per condition. **B**. Principle coordinate analysis representing bacterial (left) and fungal (right) larval gut beta diversity as determined by Bray-Curtis dissimilarity. Three biological replicates per group. Y.F2 = Yuba F2 treated with wild-sourced frass extract. Y.F6 = Yuba F6 treated with F5 captive frass extract. DOM = Domestic strain treated with domestic frass extract. **C**. The top 10 most abundant bacterial OTUs in the gut of Domestic larvae treated with domestic frass are listed to the right and aligned with the relative abundance of these OTUs in Y.F2 larvae treated with wild-sourced frass extract and Y.F6 larvae treated with F5 captive frass extract (blue = bacteria). The three most abundant fungal OTUs (across all conditions) are shown at the bottom (magenta = fungi). **D**. Volcano plot displaying fold-change in abundance (x-axis) and statistical significance (y-axis) for gut bacterial OTUs in Y.F6 larvae treated with F5 captive frass extract divided by abundance in Y.F2 larvae treated with wild-sourced frass extract. Only those OTUs with fold-change greater than or equal to 2 (log_2_ ≥ 1.0) and adjusted p-values less than 0.05 (-log10(padj) > 1.30) are labeled by name. OTUs of interest (from the top 10 list in panel C) that did not meet the adjusted p-value cutoff are labeled in italics and “n.s.” for not significant.

The attenuation of growth promotion that occurs with captive frass might provide clues to which microbes are competent to support growth on rabbit manure, and which microbes fail to support growth. To search for such clues, we compared the gut microbiomes of Y.F2 inoculated with wild-sourced frass extract, Y.F6 inoculated with F5 captive frass extract, and Domestic strain inoculated with domestic frass extract, all when reared on lab diet. Beta diversity analysis showed that F5 captive frass established bacterial gut microbiomes that were more similar to domestic frass-associated gut microbiomes and less similar to wild frass-associated gut microbiomes (Fig. 5B). A similar outcome was observed for fungal gut microbiomes. Notably, six genera (*Dysgonomonas*, *Providencia*, *Morganella*, *Enterococcus*, *Klebsiella*, and *Actinomyces*) that are commonly found in the gut of domestic BSFL^21,42^, increased in abundance over the course of captive rearing and were similarly represented in the Y.F6 (late captive frass-treated) and Domestic guts (Fig. 5C). There were very few fungal OTUs detected in lab diet-reared frass and guts (in contrast to the diversity found in rabbit manure-reared frass and guts) and shifts during captive rearing were largely binary. Wild-sourced frass treated Y.F2 guts were almost entirely populated with *Pichia*. The *Pichia* OTU is lost in Y.F6 and Domestic guts, with *Suhomyces* becoming the sole fungal OTU. There is also a shift in the total number of ITS reads that map to fungi, with the “other/unknown” category increasing in Y.F6 and Domestic samples (Fig. 5C). As previously described^43,44^, these sequences are almost entirely non-fungal reads (mainly mapping to plants present in the lab diet) and their abundance is inversely correlated with the abundance of fungal-mapping sequences. This result suggests that wild-sourced frass inoculation supports higher levels of fungal growth (nearly all *Pichia*) versus later generation captive frass or domestic frass inoculation. *Scedosporium* and *Petriella* are only detected at very low levels (average abundance < 0.1%) across all lab diet samples (SUPPLEMENTAL FILE 2).

Comparing the gut microbiome of Y.F6 larvae treated with captive frass to Y.F2 larvae treated with wild-sourced frass confirmed significant changes in bacterial abundance, including depletion of *Rummeliibacillus* spp. and enrichment of domestication-associated bacteria in the Y.F6 samples (Fig. 5D). These results show that successive generations of captive rearing on a lab diet, even after initial seeding with wild-sourced frass, shift the BSFL gut microbiome toward a domestic profile.

### Wild-sourced frass is sufficient for growth on cattle manure

If the candidate microbes described above broadly support BSFL growth on high lignocellulose animal waste, similar outcomes should be observed with cattle manure. Naïve (absent any BSFL or frass treatment) cattle manure is populated by a different set of bacteria and fungi than naïve rabbit manure (Fig. 6A). This difference allowed us to test if wild-sourced frass extract has consistent effects on growth phenotypes and gut microbe content regardless of the naïve substrate microbiome. We tested Domestic strain growth on cattle manure and found that, similar to rabbit manure, larvae grew very poorly in the absence of frass and when inoculated with domestic frass (Fig. 6B). In contrast, larval growth was significantly increased by wild frass inoculation. Gut microbiome analysis revealed significant enrichment of *Rummeliibacillus* and *Porphyromonadaceae*, in addition to an OTU not previously associated with wild-sourced frass inoculation, *Proteiniphilum* (Fig. 6C). Fungal microbiome analysis further supported the relevance of *Scedosporium* and *Petriella*. These results show that, in spite of different naïve substrate microbiomes, the benefits of wild-sourced frass extract extend to cattle manure and are likely due to conserved effects on the BSFL gut microbiome.

**Figure 6.**
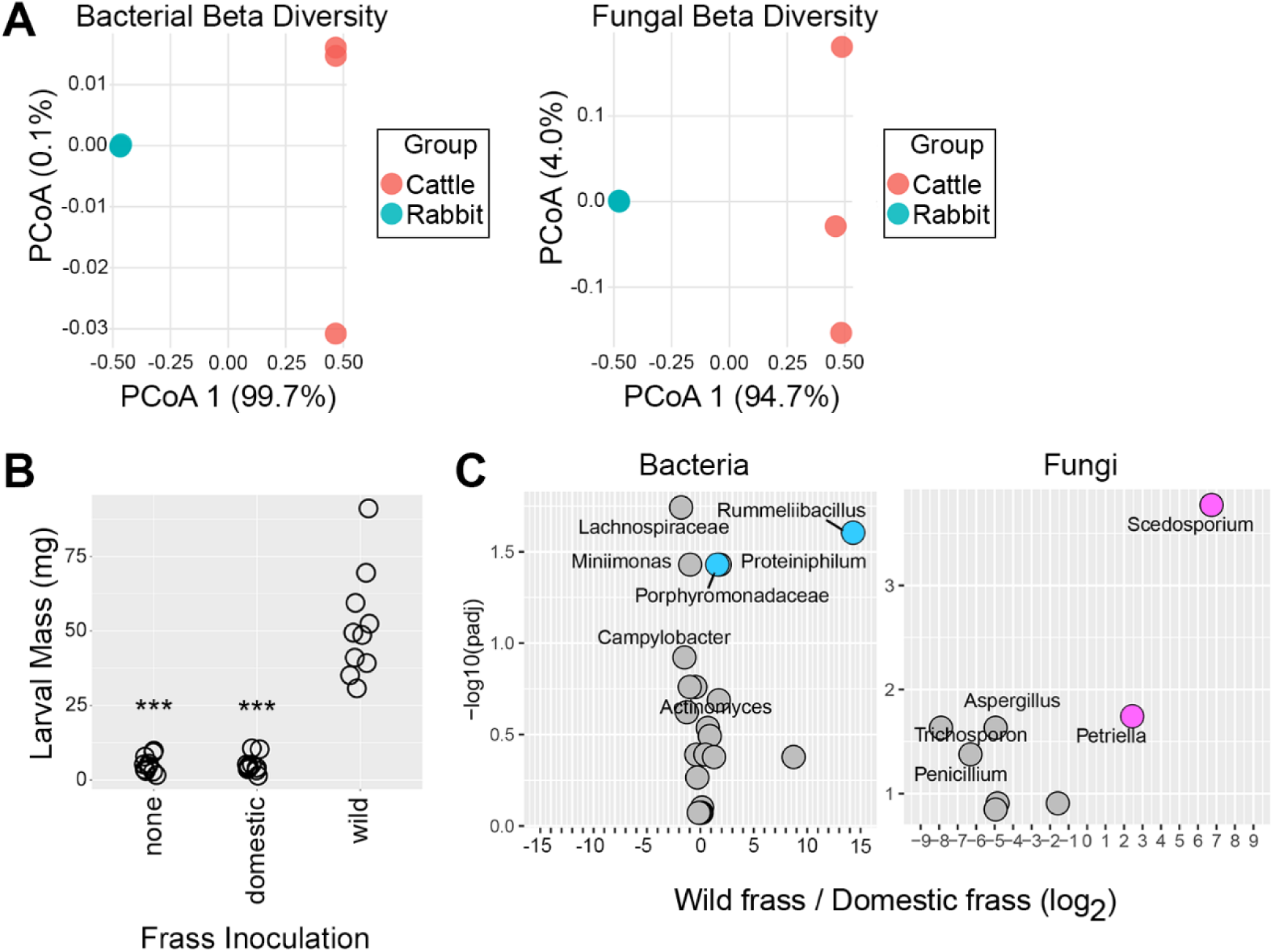
Wild-sourced frass extract from rabbit manure enables growth on cattle manure. **A**. Principle coordinate analysis representing bacterial (left) and fungal (right) larval gut beta diversity as determined by Bray-Curtis dissimilarity. Three biological replicates per group. Cattle = naïve cattle manure (no BSFL or frass treatment). Rabbit = naïve rabbit manure (no BSFL or frass treatment). **B**. Larval mass following 21 days of rearing Domestic strain on cattle manure. Frass treatments indicated on the x-axis. *** indicates p-value < 0.0001 compared to treatment with wild-sourced frass extract. n = 10 larvae per condition. **C.** Volcano plot displaying fold-change in abundance (x-axis) and statistical significance (y-axis) for gut bacterial OTUs (left) and gut fungal OTUs (right) in Domestic larvae reared on cattle manure. Fold-change is based on wild-sourced frass extract treatment divided by domestic frass extract treatment. Only those OTUs with fold-change greater than or equal to 2 (log_2_ ≥ 1.0) and adjusted p-values less than 0.05 (-log10(padj) > 1.30) are labeled by name. Bacterial OTUs of interest are labeled blue, fungal OTUs of interest are labeled magenta.

## DISCUSSION

Wild *Hermetia illucens* encounter a range of low-quality foods with difficult to obtain nutrients. These populations are under selective pressure for the holobiont (the fly plus its microbial symbionts) to evolve traits that enable bioconversion of challenging substrates. Natural selection has likely established many locally adapted wild populations of Black Soldier Flies with phenotypes of interest, such as the ability to bioconvert agricultural waste. This study was inspired by one such population and sought to determine if the desired trait (bioconversion of lignocellulose-rich animal manure) may be transferred via frass-associated microorganisms.

### Genes versus environment: frass is the primary determinant of BSFL growth

Microsatellite data showing genetic similarity among the Yuba and Domestic populations were important for contextualizing larval growth and microbiome results. Within the Yuba strain, there was no genetic drift between the P0 and F6 generations. This maintenance of genetic structure supports the conclusion that phenotypic differences among Yuba generations are more likely due to the environmental frass-associated microbiome than genes. For example, if the population had gone through a major bottleneck, differential growth of Y.F2 and Y.F6 larvae treated with captive frass could primarily be due to genetic determinants. Microsatellite data also suggest genetic similarity between Yuba and Domestic BSF. Such relatedness is expected when comparing North American BSF populations since prior studies found minimal genetic difference, even between wild and domestic strains^41,45^. However, the divergent life histories and selective pressures faced by the Yuba and Domestic strains make it possible that genetic differences exist at growth-related loci. Even if such heterogeneity exists, our data suggest that the frass extract-derived microbiome is a stronger determinant of growth. Frass effects likely intersect with host genetics^46,47^ and investigation of frass-dependent microbiome remodeling combined with high resolution genotyping of distinct lineages could prove informative.

### A small cohort of gut microbes are linked to growth on high lignocellulose manures

Through manipulation of a single variable, the frass extract, we were able to link larval growth to specific changes in gut microbiomes. These data support two general conclusions. First, host selection establishes gut microbiomes that are distinct from substrate frass microbiomes. This agrees with prior studies that found extensive differences between the relative abundance and taxonomic identities of microbes in substrate frass versus guts^16,17^. Our findings support an emerging model in which low frequency microbes of the frass community are selectively recruited and amplified (via conditions that support proliferation, eliminate competition, or both) in the gut microenvironment^23,24,28,48–51^. The second general conclusion is that changes in the gut microbiome are relevant to growth on lignocellulose-rich animal manures. This agrees with evidence that bacteria expressing lignocellulose-degrading enzymes are selectively recruited to defined BSFL gut compartments and support lignocellulose catabolism within those environments^50^, but optimal bioconversion of lignocellulosic mass likely relies on microbial activity in substrate frass and the gut^23,52^. Relative contributions of each community likely depend on the abundance of key microbes and temporal dynamics. Cattle manure experiments additionally suggest a strong influence of frass-derived microbes. Despite the fact that cattle manure and rabbit manure have very different naïve microbiomes, nearly identical OTUs were enriched in the gut of wild frass-treated larvae that grew on cattle manure. This suggests that the naïve substrate microbiome is somewhat irrelevant (a possible exception is mentioned below) and that frass seeding of naïve substrate has a dominant effect on larval growth.

Analysis of differential OTU abundance identified a small cohort that was consistently enriched in the gut of BSFL supplemented with wild-sourced frass and consistently diminished in the gut of BSFL supplemented with captive frass. Among bacteria, an OTU within the *Rummeliibacillus* genus was significantly linked to growth in all experiments. Members of *Rummeliibacillus* are gram-positive, spore-forming, rod shaped bacteria in the *Caryophanaceae* family^53^. This genus has previously been identified in BSFL gut microbiome surveys but at relatively low abundance and functional relevance was therefore not addressed^34,54–56^. In contrast, we found that *Rummeliibacillus* was the most abundant OTU in the gut of wild frass-treated Y.F2 larvae grown on rabbit manure (18.2 ± 1.5%) and the second most abundant OTU in the gut of wild frass-treated Domestic larvae grown on rabbit manure (8.2 ± 1.5%) (Fig. 3, SUPPLEMENTAL FILE 1). *Rummeliibacillus* abundance in wild frass-treated Domestic larvae fed cattle manure was 2.0 ± 0.5% (#17 in ranked abundance) (SUPPLEMENTAL FILE 1). Low abundance bacteria can still have large influences on growth phenotypes^21^ and the relatively decreased abundance of *Rummeliibacillus* in larvae fed cattle manure may indicate competition with naïve cattle manure microbes that are absent from rabbit manure. There are several indications that *Rummeliibacillus* are involved in lignocellulose breakdown. A draft *Rummeliibacillus* species genome includes genes involved in lignocellulose metabolism^57^. Growth-promoting and lignocellulose-degrading effects of *Rummeliibacillus* bacteria have been described in fish; *Rummeliibacillus* isolated from the intestine of flounders or tilapia promoted growth, feed conversion, immune function and intestinal cellulase and xylanase activity when provided as a probiotic^58,59^. A clue to another possible BSFL-related function comes from the identification of *Rummeliibacillus* in larvae fed a 70% keratin diet^56^. Animal fur is almost entirely composed of keratin protein and is highly resistant to degradation^60^. Rabbit manure used in this study contained fur and it is possible that the growth benefit associated with *Rummeliibacillus* came, at least in part, from breakdown of fur.

The other OTUs linked to growth on rabbit manure are known to function in lignocellulose digestion. Members of the genus *Bacillus* and the related genus *Paenibacillus* are important contributors to nutrient cycling in soil and manure environments and express a broad range of cellulases^61^. Members of the family *Porphyromonadaceae* perform a broad range of ecosystem functions related to metabolism of plant fibers and are particularly abundant in the gut of wood-feeding termites^62^.

The fungi enriched in the gut of wild frass-treated BSFL, *Scedosporium* and *Petriella*, are also involved in lignocellulose degradation. *Scedosporium* species are broadly implicated in wood decay^63^ and have been found in *Nasutermites* termite workers^64^ as well as the larval gut of multiple species of wood-feeding beetle^65,66^. *Scedosporium* isolated from beetle larvae in Costa Rica had cellulase, *β*-D-xylanase, and *β*-D-glucanase activity^66^. Like *Scedosporium*, *Petriella* is a member of the *Microascacae* family and is often found enriched in manure or compost^67,68^. *Petriella setifera* is known to secrete cellulase and to degrade lignin^69,70^. *Petriella* are also able to metabolize byproducts of lignocellulose degradation^71^ and therefore might contribute to carbon cycling in the BSFL microecosystem. We are unaware of previous descriptions of *Petriella* species as residents of insect guts. This OTU was also detected in substrate frass and growth-promoting effects might derive from activity in the frass and the gut.

Each of the above microorganisms are significantly linked to growth on livestock manure but mechanistic proof will require identification at the species level and functional validation via inoculation of pure cultures. Prior evidence suggests that lignocellulose metabolism in BSFL relies on multiple enzymes provided by a consortium of bacteria^22^. Our results suggest that synergistic functions of a small cohort of bacteria and fungi may optimize lignocellulose breakdown in manures.

### Growth-promoting effects of wild-sourced frass, and associated microbes, are lost during domestication

Wild-sourced frass supported growth on lab diet even though the resulting gut microbiome was significantly different from that of domestic frass-treated larvae. This suggests that the necessary metabolic functions for growth on a rich diet may be provided by a range of bacteria and fungi. This agrees with the notion that, as stated by Gligorescu et al., “the gut microbiome is functionally stable but taxonomically flexible”^42^. Taxonomic flexibility does not appear to extend to lignocellulose-rich manures, however, since specific taxonomic changes strongly correlate with larval growth.

Captive rearing of the Yuba strain provided a view into how standard BSF husbandry may bias gut microbiomes. F1 captive frass was somewhat competent to support growth on rabbit manure but not to the degree induced by wild-sourced frass. Correspondingly, the abundance of *Rummeliibacillus*, *Scedosporium*, and *Petriella* diminished compared to wild frass-treated larvae. Sequential dilution of wild-sourced frass contributions ultimately resulted in a loss of competence to support growth on rabbit manure and a corresponding shift toward bacteria commonly identified in the domestic BSFL gut. Gut microbiome changes have been shown to occur rapidly and persist across generations upon transition to a low-quality diet^72^. Even though the wild frass-associated gut microbiome initially supported growth on a lab diet, microbes that were unique to wild frass-treated guts (e.g. *Rummeliibacillus*) were lost during captive rearing. This shift may occur because nutrient-rich diets alter the gut physiology in ways that favor the domestic cohort of bacteria or there may be energetic and nutritional costs that disfavor maintenance of *Rummeliibacillus* and related microbes in the absence of selective pressures. The loss of gut fungal diversity in BSFL reared on lab diet compared to rabbit manure and the overall decreased abundance of fungi in late captive frass-treated larval guts agrees with prior findings that fiber-rich frass supports a more complex fungal microbiome^73^ and that fungal abundance is low in nutrient-rich diets^25^. Regular inoculation of wild-sourced frass extract, or repeated transfers of total frass fractions, may be necessary to maintain a gut microbiome of interest. This approach has been used to establish and even augment BSFL bioconversion of lignocellulose-rich maize straw; frass transfers drove taxonomic divergence between the gut and frass at each generation, ultimately favoring *Enterococcus* and *Actinobacteria* in the gut versus *Gammaproteobacteria* and *Dysgonomonas* in the frass, and these changes correlated with increased lignocellulose degradation in each environment^23^.

### Mechanisms of frass-mediated gut microbiome remodeling and potential applications

It remains to be determined if the benefits of frass extract inoculation are relevant to the functions of frass and frass transfer in wild populations of BSF. BSF females prefer to lay eggs in frass compared to naïve substrate^74,75^ and BSF larvae prefer substrate colonized by conspecifics over naïve substrate^76^. These discriminating behaviors may have evolved so that larvae hatch onto or seek out substrate primed with beneficial microorganisms able to colonize their gut. Frass-associated microorganisms might travel along with larvae; BSFL will emigrate from habitats with unfavorable conditions (such as high heat or intense light^77,78^) and potentially seed new substrates with microbes on their surface and in their feces. The 5% frass volume used in our experiments is likely greater than what might be transferred in the wild, although large concentrations of BSFL could produce considerable amounts of feces over time; a single late-stage larvae produces 7.8 mg of feces over a 24 hour period^79^. Inoculation of 5% frass is likely within the range of frass carryover that occurs in habitats where naïve substrate is routinely added to existing populations of BSFL, such as compost bins and manure piles.

For research and industrial applications, it may be beneficial to replace frass extract with microbial cultures. This type of probiotic treatment has previously been applied to BSFL^48,80^ and here we provide a list of promising novel probiotic candidates. A relevant finding is that Domestic strain larvae reared on lab diet for many generations benefited from a single application of wild-sourced frass extract at the time of transfer to naïve rabbit manure. This suggests that frass extracts or probiotic doses do not need to be applied over multiple generations and can be used on an ad hoc basis. Such applications may enable flexible waste bioconversion operations in which BSFL could efficiently be switched between different waste substrates by providing an appropriate reprogramming dose of frass extract.

### Conclusions

Natural selection may give rise to locally adapted populations of *Hermetia illucens* whose larvae are able to grow on unique substrates. We investigated the growth-promoting properties of environmental frass sourced from one such population: BSFL able to thrive on high-lignocellulose rabbit manure. Wild-sourced frass extract was necessary and sufficient to enable growth on lignocellulose-rich livestock manures. A small cohort of microorganisms was enriched in the gut of wild frass-treated BSFL that grew on manure and depleted in the gut of captive frass-treated BSFL that failed to grow on manure. This cohort was consistently centered around *Rummeliibacillus* bacteria and *Scedosporium* fungi, suggesting a bacterial-fungal synergy supporting lignocellulose catabolism and bioconversion. This work demonstrates the value of incorporating BSF microecosystems from wild environments into studies of BSFL biology and industrial potential.

## MATERIALS AND METHODS

### Field Collections and Laboratory Husbandry

Rabbit manure containing a high concentration of BSFL at all stages of development and separate samples of naïve rabbit manure with no observable BSFL were collected from a rabbit farm in Yuba County, CA, USA. Samples were collected from neighboring open-air trays. The trays were positioned below rabbit cages and captured a mixture of rabbit feces, urine and fur. The rabbit manure containing BSFL was transferred to the laboratory and placed in cross-stacked large plastic bins inside two individual 60 cm x 60 cm x 120 cm nylon netting population cages (6E620 insect rearing cage, BugDorm Store, MegaView Science Co., Ltd., Taiwan). This wild-sourced rabbit manure + BSFL served as the source of Yuba P0 adults as well as the wild-sourced frass extract used in subsequent experiments. Founding P0 adults mated and laid eggs in the substrate (eggs were also laid in collection bins that were removed to establish the captive bred population, described below); at that point the wild-sourced frass contained a mixture of Yuba P0 and F1 larvae and naïve rabbit manure from the farm source was added to continue supporting larval development (∼20% of the initial volume was added). This wild-sourced frass microenvironment was maintained for 60 days. Prior to any use, the BSFL-free rabbit manure was examined under a microscope to confirm the absence of any insects and homogenized by mixing in one large batch. Small batches were stored at −20°C; these were thawed as needed and served as naïve rabbit manure in subsequent experiments.

Captive rearing of the Yuba strain was performed using a scheme in which frass from one generation seeded the naïve lab diet substate of the subsequent generation. This occurred because larvae of the parent generation freely accessed containers of lab diet in which eggs of the subsequent generation were collected. Beginning with the Yuba P0 founding sample, 8 open collection bins (4.4 inch diameter, 500 ml Nalgene polypropylene jars, Thermo Scientific, USA) filled with 350 ml lab diet were deeply embedded in the wild-sourced frass. Lab diet was 31.5% corn meal, 31.5% wheat bran, 25% glucose, 12% yeast extract, with the dry mixture suspended in an equal volume of sterile water. This preparation was used in collection bins, Domestic population cages and growth assays. Within the P0 population cages, larvae freely accessed these collection bins, seeding the lab diet with frass carried on their surface and with larval excrement. P0 adults that emerged within these population cages laid eggs directly in the lab diet and within strips of corrugated cardboard set into the collection bins. Eggs were either removed for use in growth assays (described below) or allowed to hatch within the collection bins. One week after the first egg clutches were observed, collection bins were removed from the population cage and any prior generation BSFL (distinguished by size) were manually cleared so that only eggs and recently hatched larvae of the next generation remained. Collection bin contents were combined with 5 liters of naïve lab diet and split into two new population cages. Eight fresh collection bins filled with lab diet were embedded in this substrate and the cycle was repeated. Domestic strain BSF (obtained from Symton Black Soldier Fly, College Station, TX, USA) were reared under similar conditions, with frass from parental generations transferred into the substrate of subsequent generations via larval activity. Population cages were kept in a climate-controlled environment, using standard BSF husbandry conditions^81^ with full spectrum (380 nm – 780 nm) LED light on a 12-hour cycle and continuous blue 460 nm LED light.

BSFL-free cattle manure, < 6 hours old, was collected from open rangeland at a cow-calf operation in Mariposa County, CA, USA. The manure was transferred to the laboratory and kept at 22°C for one day prior to use in growth assays.

### Growth assays and frass extract inoculation

BSF eggs were harvested from collection bins, rinsed with sterile water on mesh screens to remove frass, then transferred to moistened tissue paper in small petri plates kept at 27°C until eye spots were observed. 30-40 eye spot-stage eggs were transferred to culture bottles containing 50 ml of substrate (naïve rabbit manure, lab diet, or naïve cattle manure). Prior to addition of eggs, the substrate was thoroughly mixed with 2.5 ml of the indicated frass extract. Frass extract was prepared under aseptic conditions by wrapping a sample of total frass in sterile 660 μm aperture mesh and squeezing the liquid fraction into sterile collection tubes. Sterile-filtered frass was prepared by passing liquid frass extract through a 0.2 μm filter.

Growth assays were performed in duplicate, and bottles were kept at 27°C. Larval mass was measured 21 days after larval hatching and five larvae were randomly selected from each bottle to obtain ten total measurements. The complete digestive tract was dissected from three of the randomly selected larvae per condition for DNA extraction.

### Microsatellite genotyping and population genetics

Previously described multiplex PCR reactions (Hi_1 and Hi_2 primer mixes) were used^36^. Fragment analysis was performed using an Applied Biosystems 3730XL DNA Analyzer and Peak Scanner 2 software. All population genetic analyses were performed using the PopGenReport package in R.

### Microbiome sequencing

Frass and dissected BSFL gut DNA for microbiome sequencing was prepared using the DNeasy PowerLyzer PowerSoil Kit (Qiagen). Library construction, sequencing, processing, read mapping and OTU quantification was performed by ZymoBIOMICS Targeted Sequencing Service. Bacterial 16S ribosomal RNA gene sequencing (V3_V4 region) was performed using the *Quick*-16S NGS Library Prep Kit (Zymo Research) and the same kit was used for fungal sequencing but with custom ITS2 primers. Libraries were sequenced on an Illumina Nextseq with a P1 reagent kit (600 cycles). The sequencing was performed with 30% PhiX spike-in.

Unique amplicon sequences were inferred from raw reads using the DADA2 pipeline^82^. Potential sequencing errors and chimeric sequences were also removed with the Dada2 pipeline. Taxonomy assignment was performed using Uclust from Qiime v.1.9.1.

### Statistical analysis

Statistical analyses were performed using R. The limma package was used to identify significant differences in OTU abundance. The inputs for limma included all OTUs with average abundance ≥ 1.0% in either condition. Bray-Curtis dissimilarity was calculated using the Vegan Community Ecology package. Significant differences in larval growth data were identified by ANOVA and TukeyHSD post-hoc tests.

## ACKNOWLEDGEMENTS

This project was funded by grants from the UC Merced Senate and the University of California Climate Action Seed Fund to M.C. We are grateful to Gordon Bennett and Ryan Torres for helpful discussions and advice during the planning and execution of this project.

M.C. acquired funding for the study. M.C. and A.P. established the experimental design and methods. A.P., R.S., K.T., and M.C. performed the experiments and collected data. M.C. carried out the bioinformatic and statistical analyses. M.C. wrote the first draft of the manuscript and feedback from all authors was incorporated. All authors read and approved the final manuscript.

